# Functional analyses of the CIF1-CIF2 complex in *Trypanosoma brucei* identify the structural motifs required for complex formation and cytokinesis

**DOI:** 10.1101/196238

**Authors:** Huiqing Hu, Paul Majneri, Dielan Li, Yasuhiro Kurasawa, Tai An, Gang Dong, Ziyin Li

## Abstract

Cytokinesis in trypanosome occurs uni-directionally along the longitudinal axis from the cell anterior towards the cell posterior and requires a trypanosome-specific CIF1-CIF2 protein complex. However, little is known about the contribution of the structural motifs in CIF1 and CIF2 to complex assembly and cytokinesis. Here, we demonstrated that the two zinc-finger motifs but not the coiled-coil motif in CIF1 are required for interaction with the EF-hand motifs in CIF2. We further showed that localization of CIF1 depends on the coiled-coil motif and the first zinc-finger motif and that localization of CIF2 depends on the EF-hand motifs. Deletion of the coiled-coil motif and mutation of either zinc-finger motifs in CIF1 disrupted cytokinesis. Further, mutation of either zinc-finger motif in CIF1 mis-localized CIF2 to the cytosol and destabilized CIF2, whereas deletion of the coiled-coil motif in CIF1 spread CIF2 over to the new flagellum attachment zone and stabilized CIF2. Together, these results uncovered the requirement of the coiled-coil motif and zinc-finger motifs for CIF1 function in cytokinesis and for CIF2 localization and stability, providing structural insights into the functional interplay between the two cytokinesis regulators.

---

*Trypanosoma brucei*, an early branching parasitic protozoan, causes lethal infections in humans and animals in sub-Saharan Africa. The parasite has a complex life cycle by alternating between insect vectors and mammalian hosts, and proliferates through binary cell fission in the midgut of insects and the bloodstream of mammals. Cell division occurs uni-directionally along the longitudinal axis from the cell anterior to the cell posterior, as visualized by time-lapse video microscopy of dividing cells (1), and the division plane appears to be determined by the length of the new flagellum and a proteinaceous filament termed flagellum attachment zone (FAZ) (2,3). The anterior tip of the newly assembled FAZ filament constitutes the site from which cytokinesis cleavage furrow starts to ingress towards the posterior cell end along the division fold, which is formed through membrane invagination by unknown mechanisms (4,5). Due to the apparent lack of involvement of actin in cytokinesis (6), trypanosome likely does not assemble an actomyosin contractile ring, the evolutionarily conserved machinery for cleavage furrow ingression found in amoebas, fungi, and animals (7,8).

The molecular mechanism for the uni-directional, longitudinal binary cell fission in trypanosome remains poorly understood. Based on the observed cell cycle defects through RNAi-mediated gene ablation, many proteins have been reported to be involved in cytokinesis, which localize to various subcellular structures and possess diverse biochemical functions (5,9-23).However, many of these proteins localize to the cytosol, raising the question of whether they play a direct role in cytokinesis. Several proteins, including the Polo-like kinase TbPLK (10),the chromosomal passenger complex (CPC) composed of the Aurora B kinase TbAUK1, TbCPC1, and TbCPC2 (13), and two trypanosome-specific proteins, CIF1 (5), which was named TOEFAZ1 in another study (20), and CIF2 (21), localize to the anterior tip of the new FAZ at different stages of the cell cycle and are involved in cytokinesis initiation. The CPC and CIF1 additionally localize to the cleavage furrow during cytokinesis (1,5,13), but whether they play any role in cleavage furrow ingression, membrane remodeling at the furrow, or daughter cell abscission remains to be investigated.

We recently uncovered a signaling cascade that acts at the anterior tip of the new FAZ to promote cytokinesis in trypanosome (5,21). This signaling cascade is composed of two evolutionarily conserved protein kinases, TbPLK and TbAUK1, and two trypanosome-specific proteins, CIF1 and CIF2, that form a complex during the S phase of the cell cycle (21). CIF1 and CIF2 are interdependent for their stability, and CIF2 protein abundance appears to be tightly regulated, as CIF2 protein disappears from the new FAZ tip after S phase and overexpression of CIF2 causes a cytokinesis defect (21). Despite their essential physiological roles, the biochemical function of CIF1 and CIF2 is still unknown. In this report, we carried out functional analyses of the structural motifs in CIF1 and CIF2 to investigate their requirement for CIF1-CIF2 complex formation and cytokinesis initiation, using yeast two-hybrid assay, size exclusion chromatography, co-immunoprecipitation, and RNAi complementation. Moreover, the requirement of CIF1 structural motifs for CIF2 localization and stability was also investigated.

## Results

### Identification of the structural motifs required for CIF1-CIF2 complex formation

The CIF1 protein sequence retrieved from *T. brucei* 927 strain genome database is 793 amino acids in length and has a calculated molecular mass of 89.6 kDa. However, when the *CIF1* gene sequence was PCR amplified from the procyclic 427 strain and sequenced, we detected a 33-nt insertion between nucleotides 534-535, which encoded an 11-aa sequence RKRKQREGEEE, resulting in a 804-aa protein. Sequencing of multiple PCR products from both the Li and the Dong laboratories confirmed this 33-nt insertion, indicating that *CIF1* gene sequence in *T. brucei* 927 strain genome database missed the 33-nt sequence during sequence annotation, likely because the 11-aa sequence encoded by the missing 33-nt sequence belongs to one of the six repetitive sequences (Figs. 1A, S1 and S2). The CIF1 homolog in *T. brucei gambiense* genome (Tbg972.11.17730) is also 804-aa in length and contains the 11-aa sequence (RKRKQREEEEE) that is only one residue different from that in *T. brucei* 927 strain (Figs. S1 and S2). However, the CIF1 homolog in *T. brucei* 427 strain genome database (Tb427tmp.01.7450) contains a 13-aa deletion and a frame shift (Figs. S1 and S2), likely due to erroneous sequence annotations. Therefore, CIF1 protein is 804-aa in length and has a calculated molecular mass of 91.0 kDa. Program COILS (24) predicted a coiled-coil motif in the N-terminal portion between a.a. 121-271, and homology modeling using the SWISS-MODEL software (25) detected two CCHC (Cysteine-Cysteine-Histidine-Cysteine)-type zinc-finger (ZnF) motifs at the C-terminus of CIF1 (Fig. 1A). Within the coiled-coil motif, there are either three 20-aa repeats or six 10-aa repeats (Fig. 1A). Both zinc-finger motifs, ZnF1 and ZnF2, share similar folds and contain the conserved zinc ion-coordinating CCHC residues (Fig. 1A). CIF2 contains a calmodulin-like domain at the N-terminus, which is composed of four EF-hand motifs, EF1-EF4 (Fig. 1B). However, all four EF-hand motifs lack at least one conserved calcium-binding residue (Fig. 1B, highlighted in red), raising the question of whether the calmodulin-like domain in CIF2 is capable of binding to calcium ions.

**Figure 1.**
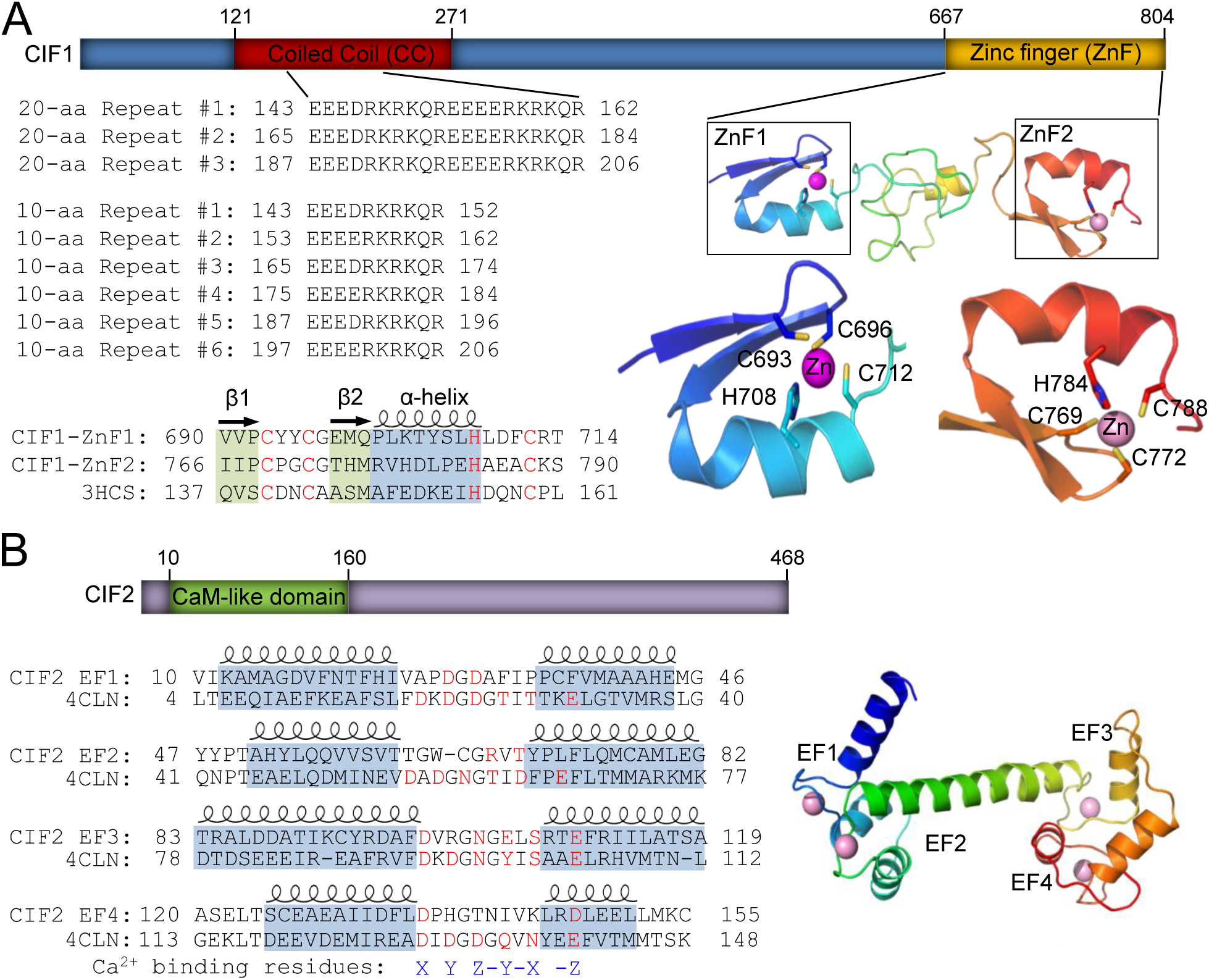
Identification of structural motifs in CIF1 and CIF2 by homology modeling. **(A)**. Illustration of the conserved domains in CIF1. The coiled-coil (CC) motif located at the N-terminal portion of CIF1 contains a repetitive sequence composed of either three 20-aa repeats or six 10-aa repeats. The two CCHC-type zinc-finger (ZnF) motifs, ZnF1 and ZnF2, share similar folds and are predicted to be able to coordinate a zinc ion by the conserved CCHC residues (highlighted in red). Zinc-finger motifs were modeled using the crystal structure of TRAF6 (PDB code: 3HCS) as the template. (**B**). Illustration of the conserved domain in CIF2. The predicted calmodulin-like domain at the N-terminal portion of CIF2 possesses four EF-hand motifs, which is modeled using the crystal structure of *Drosophila* calmodulin as the template (PDB code: 4CLN). Note that all four EF-hand motifs lack one or more conserved residues (highlighted in red) required for coordinating the calcium ion.

To identify the structural motifs that are required for CIF1-CIF2 interaction, we first carried out yeast two-hybrid assays. Three CIF1 mutants, CIF1-ΔCC, CIF1-ZnF1^mut^, and CIF1-ZnF2^mut^, were generated by deleting the coiled-coil motif (a.a. 121-271) or by mutating the four zinc-coordinating residues (Cysteine, Cysteine, Histidine, and Cysteine) of ZnF1 and ZnF2 to alanine (Fig. 2A). Four CIF2 mutants, CIF2-ΔEF1, CIF2-ΔEF1-2, CIF2-ΔEF1-3, and CIF2-ΔEF1-4, were generated by deleting one, two, three, or four EF-hand motifs (Fig. 2A). Yeast two-hybrid assays showed that CIF2, but not CIF1, interacted with itself (Fig. 2B). Deletion of EF1 disrupted CIF2 self-interaction under high stringency (4DO, quadruple drop-out) condition (Fig. 2B), indicating that in yeast CIF2 formed a dimer or a multimer through the EF-hand motifs.

**Figure 2.**
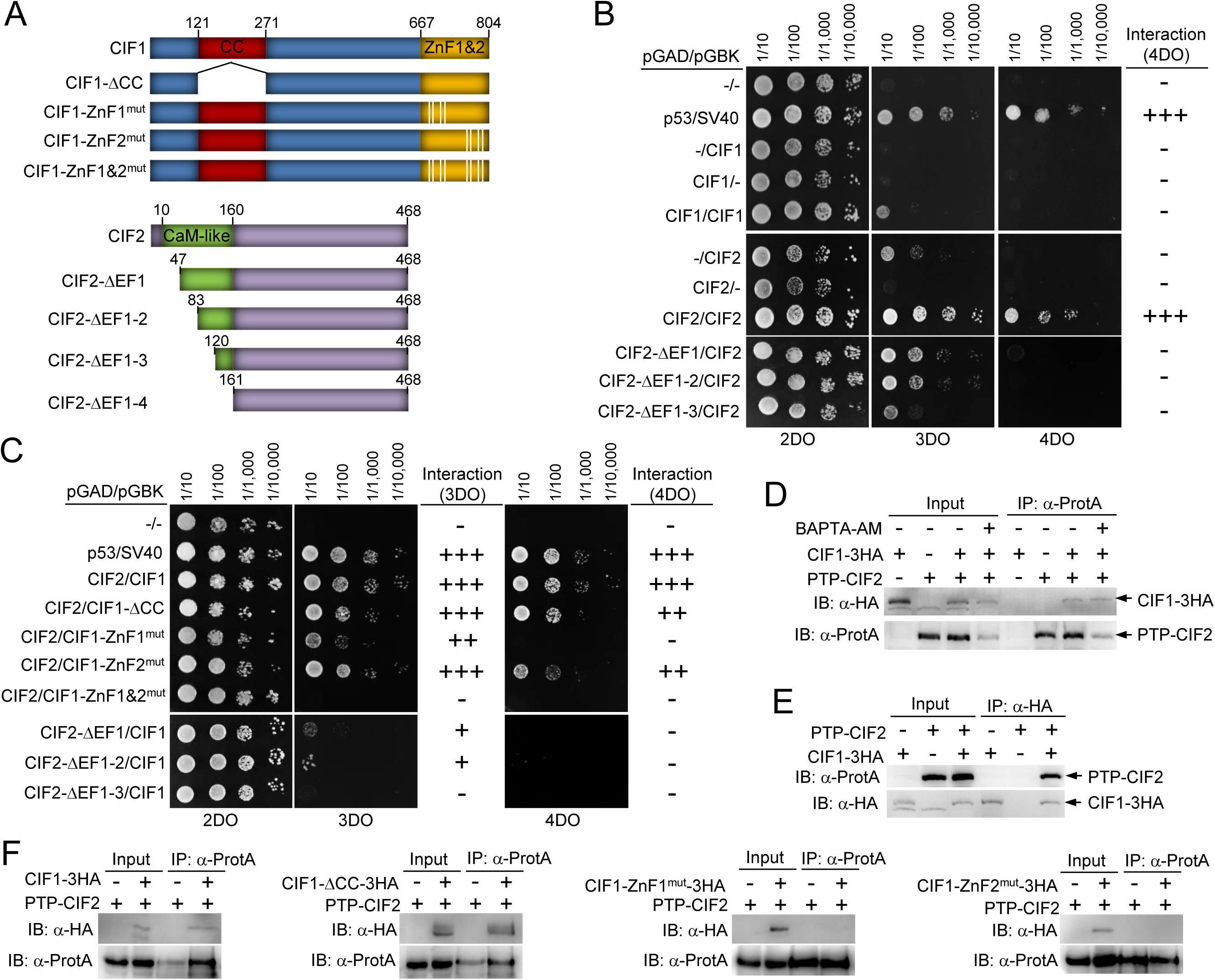
Identification of the structural motifs involved in CIF1-CIF2 interaction. (**A**). Schematic illustration of wild-type and mutant CIF1 and CIF2. CIF1-ΔCC, CIF1 coiled-coil motif deletion mutant; CIF1-ZnF1^mut^, CIF1 zinc-finger 1 mutant generated by mutating Cys693, Cys696, His708, and Cys712 to alanine residue; CIF1-ZnF2^mut^, CIF1 zinc-finger 2 mutant generated by mutating Cys769, Cys772, His784, and Cys788 to alanine; CIF2-ΔEF1, CIF2 EF-hand motif #1 deletion mutant by deleting a.a. 1-46; CIF2-ΔEF1-2, CIF2 EF-hand motifs #1-2 deletion mutant by deleting a.a. 1-82; CIF2-ΔEF1-3, CIF2 EF-hand motifs #1-3 deletion mutant by deleting a.a. 1-119; CIF2-ΔEF1-4, CIF2 EF-hand motifs #1-4 deletion mutant by deleting a.a. 1-160. (**B**). Yeast two-hybrid assays to test the potential self-interaction of CIF1 and CIF2. 2DO medium (SD/-Leu/-Trp/) selects the presence of both the pGAD and pGBK plasmids. 3DO medium (SD/-Leu/-Trp/-His/) detects protein-protein interaction under medium stringency conditions, and 4DO medium (SD/-Leu/-Trp/-His/-Ade) detects protein-protein interaction under high stringency conditions. Note that due to the self-activation of CIF2 on 3DO medium, protein-protein interactions were only scored based on the 4DO medium. (**C**). Yeast two-hybrid assays to test the potential interaction between CIF1 and CIF2 and the requirement of structural motifs for CIF1-CIF2 interaction. (**D**). Co-immunoprecipitation to test the interaction between PTP-tagged CIF2 and 3HA-tagged CIF1 in the absence or presence of the calcium chelator BAPTA-AM. Immunoprecipitation was carried out by pulling down PTP-CIF2 with IgG beads, and the co-immunoprecipiated CIF1-3HA and PTP-CIF2 were detected by anti-HA mAb and anti-Protein A (α-ProtA) pAb, respectively. Cells expressing PTP-CIF2 alone and CIF1-3HA alone were included as negative controls. (**E**). Reciprocal immunoprecipitation to test the interaction between CIF1-3HA and PTP-CIF2. Immunoprecipitation was carried out by pulling down CIF1-3HA with anti-HA agarose beads, and co-immunoprecipitated PTP-CIF2 and CIF1-3HA were detected by anti-Protein A pAb and anti-HA mAb. Cells expressing CIF1-3HA alone and PTP-CIF2 alone were included as negative controls. (**F**). Co-immunoprecipitation to test the potential interaction between CIF2 and CIF1 mutants. Wild-type and mutant CIF1 proteins were tagged with a triple HA epitope and ectopically expressed in CIF1-3’UTR RNAi cell line, and CIF2 was tagged with a PTP epitope from its endogenous locus in the CIF1-3’UTR RNAi cells expressing wild-type and mutant CIF1 proteins. PTP-CIF2 was immunoprecipitated by IgG beads, and the immunoprecipitated proteins were immunoblotted with anti-HA mAb to detect 3HA-tagged wild-type and mutant CIF1 and with anti-Protein A pAb to detect PTP-CIF2.

Interaction between CIF1 and CIF2 was detected in yeast under both medium stringency (3DO, triple drop-out) and high stringency conditions (4DO, quadruple drop-out) (Fig. 2C), indicating that the two proteins form a tight complex. Deletion of the coiled-coil motif did not affect CIF1-CIF2 interaction under both medium and high stringency conditions (Fig. 2C), indicating that the coiled-coil motif is not involved in CIF1-CIF2 interaction. Under medium stringency condition, mutation of ZnF1 significantly weakened CIF1-CIF2 interaction, whereas mutation of ZnF2 did not affect the interaction between CIF1 and CIF2 (Fig. 2C). However, mutation of both ZnF1 and ZnF2 completely disrupted CIF1-CIF2 interaction (Fig. 2C). Under high stringency condition, mutation of ZnF1 completely disrupted interaction, and mutation of ZnF2 significantly weakened the interaction (Fig. 2C). These results suggest that both ZnF1 and ZnF2 are involved in CIF1-CIF2 interaction in yeast, albeit to different extents. We next tested whether the EF-hand motifs are involved in CIF1-CIF2 interaction. Deletion of the EF1 and EF2 significantly weakened the interaction between CIF1 and CIF2 under medium stringency condition, and completely disrupted CIF1-CIF2 interaction under high stringency condition (Fig. 2C). Further deletion of EF3 completely disrupted CIF1-CIF2 interaction (Fig. 2C). These results indicated that the four EF-hand motifs as a whole are involved in CIF1-CIF2 interaction.

To corroborate the yeast two-hybrid results, we carried out co-immunoprecipitation experiments. We first tested whether CF1-CIF2 interaction requires calcium by treating the cells that co-express PTP-CIF2 and CIF1-3HA with a membrane permeable calcium chelator, BAPTA AM (1,2-bis(O-aminophenoxy)ethane-*N*,*N*,*N’,N*’ tetraacetic acid (acetoxymethyl) ester) (26,27). Immunprecipitation of CIF2 was able to pull down CIF1 in both the absence and the presence of BAPTA-AM (Fig. 2D). This further confirms that CIF2 and CIF1 form a complex *in vivo* in trypanosome, as reported previously (21), and suggests that CIF1-CIF2 interaction likely does not require calcium. To further confirm the interaction between CIF1 and CIF2, we carried out a reciprocal co-immunoprecipitation experiment, and the results showed that immunoprecipitation of CIF1 was able to pull down CIF2 from trypanosome cell lysate (Fig. 2E). Finally, to test whether mutation of the coiled-coil motif and the zinc-finger motifs in CIF1 affects its binding to CIF2, we first generated a CIF1 RNAi cell line by targeting the 3’UTR of CIF1 and then overexpressed 3HA-tagged CIF1, CIF1-ΔCC, CIF1-ZnF1^mut^, or CIF1-ZnF2^mut^ in this RNAi cell line to further generate four RNAi complementation cell lines (see below for validation of these cell lines). Subsequently, CIF2 was endogenously tagged with a PTP epitope at the C-terminus in these RNAi complementation cell lines. RNAi of CIF1 and overexpression of ectopic CIF1 and its mutants were induced by incubating the cells with tetracycline. Co-immunoprecipitation was carried out by incubating the cleared cell lysate with IgG beads to pull down CIF2-PTP and its associated proteins, which were then immunoblotted with anti-HA antibody to detect co-immunoprecipitated 3HA-tagged CIF1 and its mutants. As shown above, CIF2 was able to pull down CIF1 (Fig. 2F). CIF2 was also able to pull down CIF1-ΔCC (Fig. 2F), indicating that the coiled-coil motif is not involved in CIF1-CIF2 interaction. However, CIF2 was not able to pull down CIF1-ZnF1^mut^ and CIF1-ZnF2^mut^ (Fig. 2F), indicating that both zinc-finger motifs are required for CIF1-CIF2 interaction.

Finally, to confirm the direct interaction between the zinc-finger motifs in the C-terminal domain (CTD) of CIF1 and the EF-hand motifs in the N-terminal domain (NTD) of CIF2, we carried out *in vitro* co-elution assays by size exclusion chromatography using recombinant proteins purified from bacteria. A sample containing both non-tagged proteins was applied onto a Superdex S-200 16/60 column after co-purification on a Ni-NTA affinity column. Non-tagged CIF1-CTD and CIF2-NTD were co-eluted at approximately 85 ml and exhibited similar band intensity on the SDS-PAGE gel (Fig. 3, blue curve). To further verify that the co-elution of CIF1-CTD and CIF2-NTD was indeed due to the formation of an intermolecular complex rather than their similar molecular mass (15.2 kDa vs 17.4 kDa), we copurified CIF1-CTD and SUMO-CIF2-NTD or SUMO-CIF1-CTD and SUMO-CIF2-NTD for size exclusion chromatography. The fusion of an N-terminal SUMO tag provides an extra ∼18 kDa of molecular mass to the target proteins, allowing the separation of SUMO-tagged and non-tagged proteins at different elution volume. Size exclusion chromatography demonstrated that the mixture of non-tagged CIF1-CTD and SUMO-CIF2-NTD shifted the elution peak to approximately 77 ml (Fig. 3, red curve), whereas an elution peak at about 65 ml was obtained with both SUMO-CIF1-CTD and SUMO-CIF2-NTD (Fig. 3, green curve). SDS-PAGE analysis showed that the peak fractions in either case contained both proteins at a similar 1:1 molar ratio, as judged by the relative intensity of the protein bands normalized by their molecular mass. Overall, these results suggest that CIF1-CTD and CIF2-NTD form a stable 1:1 stoichiometric complex.

**Figure 3.**
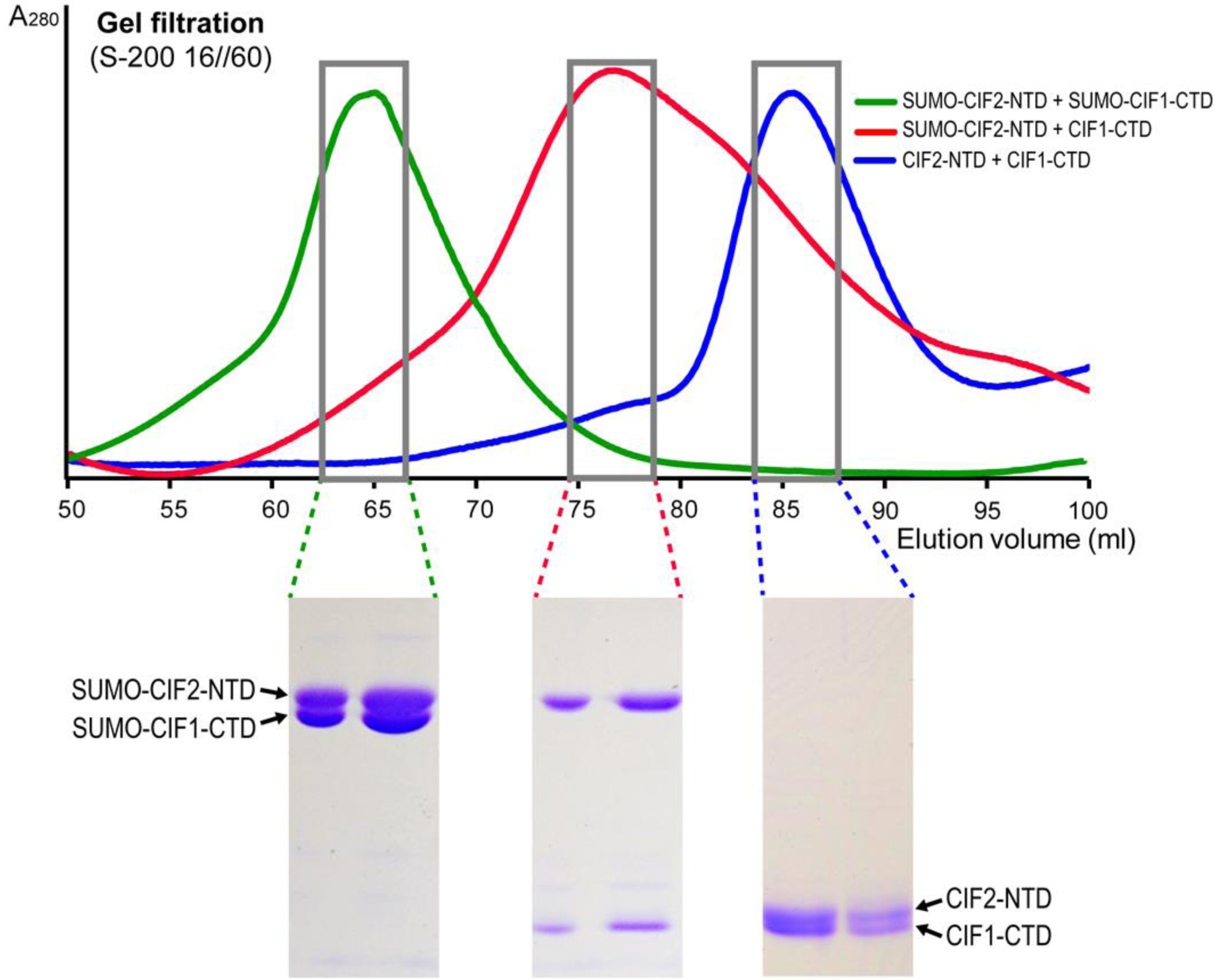
CIF1-CTD forms a 1:1 stoichiometric complex with CIF2-NTD. Size exclusion chromatography (S200 16/60) elution profiles of three different combinations of co-purified CIF1-CTD (a.a. 667-804, calculated M.M. of 15.2 kDa) and CIF2-NTD (a.a. 1-160, calculated M.M. of 17.4 kDa) are overlaid to show the shifts of the larger complexes with one or both proteins bearing a SUMO tag. Peak fractions from each elution were examined on a SDS-PAGE gel, which showed that the two proteins were co-eluted and had a 1:1 molar ratio in each case.

### Identification of the structural motifs required for CIF1 and CIF2 localization

To investigate the requirement of the structural motifs for CIF1 and CIF2 localization to the new FAZ tip, wild-type and mutant CIF1 and CIF2 proteins were ectopically expressed in the procyclic form of *T. brucei*. Expression of these proteins was induced with 0.1 μg/ml tetracycline to allow low levels of expression, which did not affect cell proliferation (data not shown). Western blotting confirmed the expression of wild-type and mutant CIF1 and CIF2 proteins (Fig. 4A, B). Immunofluorescence microscopy showed that wild-type CIF1 was localized to the new FAZ tip (Fig. 4C), demonstrating that ectopic expression of CIF1 did not affect its localization to the new FAZ tip. Deletion of the coiled-coil motif prevented CIF1 from concentrating at the new FAZ tip, but instead caused CIF1 protein to spread over to the anterior one-third length of the new FAZ in most (>75%) of the 1N2K and 2N2K cells or throughout the full length of the new FAZ in the rest of the 1N2K and 2N2K population (Fig. 4C). In most (>95%) of the 1N1K cells that possess a short new FAZ, CIF1-ΔCC was also detected at the entire new FAZ (data not shown). These results suggest that the coiled-coil motif is required for restricting CIF1 to the anterior tip of the new FAZ. CIF1-ZnF1^mut^ was detected throughout the cytosol (Fig. 4C), indicating that ZnF1 is required for CIF1 localization. CIF1-ZnF2^mut^, however, was still detectable at the new FAZ tip (Fig. 4C), suggesting that ZnF2 is not required for CIF1 localization.

**Figure 4.**
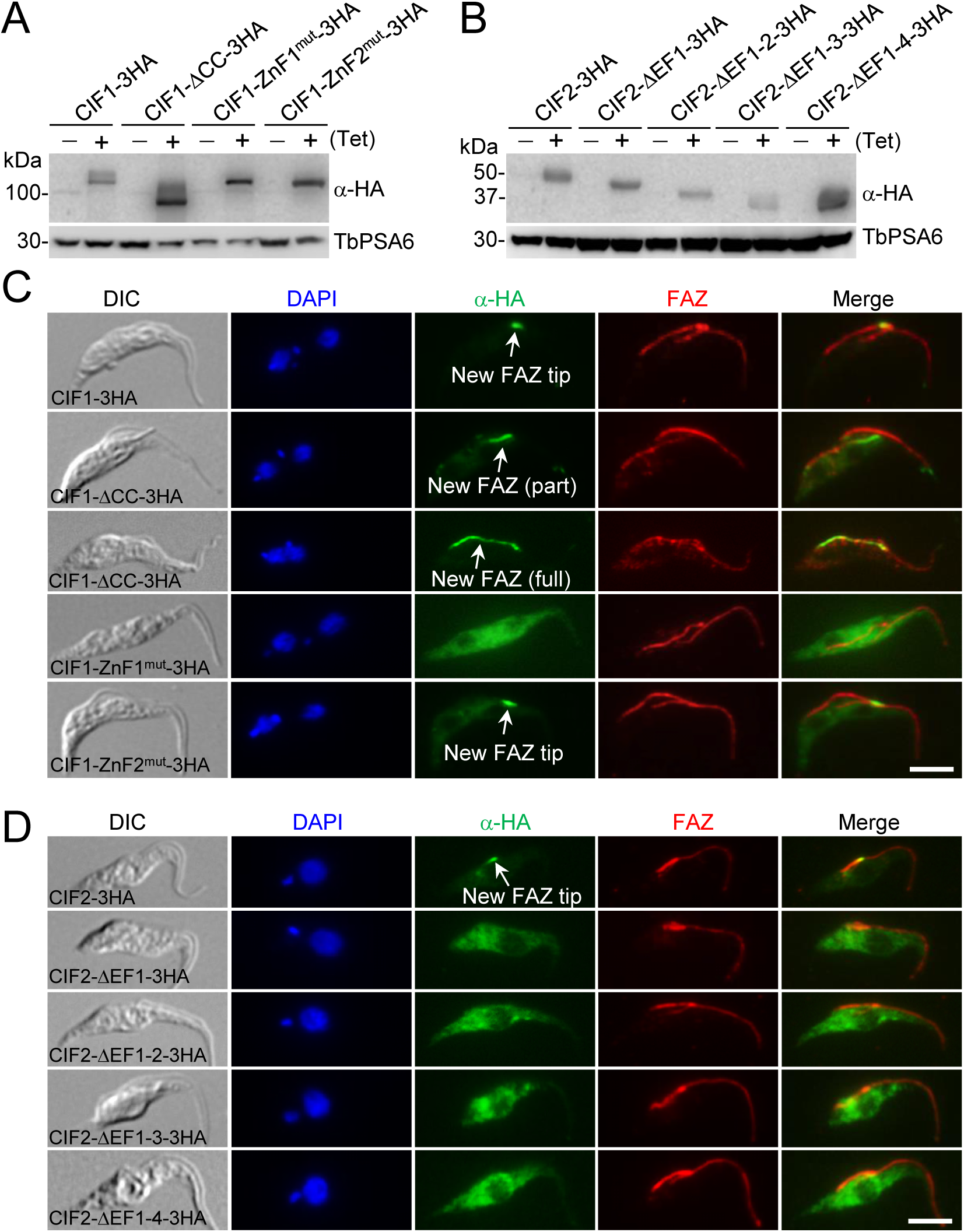
Subcellular localization of CIF1 and CIF2 mutants. (**A**, **B**). Western blotting to monitor the level of triple HA-tagged wild-type and mutant CIF1 and CIF2, which were induced with 0.1 μg/ml tetracycline for 6 h. TbPSA6 served as the loading control. (**C**). Immunofluorescence microscopy to examine the subcellular localization of wild-type and mutant CIF1. Cells were co-immunostained with FITC-conjugated anti-HA mAb to label 3HA-tagged CIF1 and its mutants and anti-CC2D pAb to label the FAZ filament. Scale bar: 5 μm. (**D**). Immunofluorescence microscopy to investigate the subcellular localization of CIF2 and its mutants. 3HA-tagged CIF2 and its mutants were detected by FITC-conjugated anti-HA mAb, and FAZ filament was labeled with anti-CC2D pAb. Scale bar: 5 μm.

When induced with 0.1 μg/ml tetracycline, wild-type CIF2 was localized to the new FAZ tip in S-phase cells (Fig. 4D), demonstrating that ectopic expression of CIF2 did not affect its localization to the new FAZ tip. In 1N2K and 2N2K cells, overexpressed CIF2 was detected at the basal body region (21), but our focus was on the S-phase 1N1K cells. Deletion of the EF1, EF1-2, EF1-3, or all four EF-hand motifs (EF1-4) of the calmodulin-like domain in CIF2, however, all disrupted CIF2 localization (Fig. 4D), demonstrating that the N-terminal EF-hand motifs are required for CIF2 localization.

### RNAi complementation experiments uncover the structural motifs in CIF1 required for CIF1 function

To examine the requirement of the structural motifs for CIF1 and CIF2 function, we carried out RNAi complementation experiments by knocking down CIF1 or CIF2 expression through RNAi against the 3’-UTR of CIF1 or CIF2 and by expressing CIF or CIF2 mutant proteins from an ectopic locus in the RNAi background. Unfortunately, this approach did not work for CIF2, as ectopic expression of CIF2 in CIF2-3’UTR RNAi cell line caused more severe growth defects than CIF2 RNAi alone (data not shown). Our previous results showed that ectopic overexpression of CIF2 and CIF2 RNAi both caused severe growth defects (21), suggesting that CIF2 level is tightly controlled. Multiple attempts to fine-tune the levels of ectopically expressed CIF2 by titrating tetracycline concentrations failed to rescue the RNAi defects. Therefore, it was not possible to test the roles of the EF-hand motifs in CIF2 function by RNAi complementation.

RNAi of CIF1 against the 3’-UTR of CIF1 was able to knock down CIF1, as demonstrated by western blotting and immunofluorescence microscopic analyses of the level of CIF1 protein that was tagged with an N-terminal PTP epitope from its endogenous locus (Fig. 5A, B). This new RNAi cell line was referred to as CIF1-3’UTR RNAi cell line to distinguish it from our previously reported CIF1 RNAi cell line that targeted the coding region of CIF1 (5). We next ectopically expressed 3HA-tagged wild-type and mutant CIF1 proteins in the CIF1-3’UTR RNAi cell line. Western blotting and immunofluorescence microscopy both confirmed the knockdown of the endogenous CIF1, which was tagged with a PTP epitope at the N-terminus, and the expression of ectopic CIF1 and CIF1 mutants, which were tagged with a C-terminal triple HA epitope (Fig. 5A, B). Using these RNAi complementation cell lines, we also investigated the localization of CIF1 and its mutants. In the CIF1-3’UTR RNAi cells, ectopically expressed CIF1 localized to the new FAZ tip, whereas CIF1-ΔCC was spread over to either the anterior one-third length of the new FAZ or the full length of the new FAZ (Fig. 5B, C). CIF1-ZnF1^mut^ was mis-localized to the cytosol, but CIF1-ZnF2^mut^ was still detectable at the new FAZ tip (Fig. 5B, C). The localization patterns of CIF1 and its mutants in CIF1-3’UTR RNAi cells are identical to that of CIF1 and its mutants that were ectopically expressed in the 29-13 strain (Fig. 4B), which further confirmed the requirement of the coiled-coil motif and ZnF1 for CIF1 localization.

**Figure 5.**
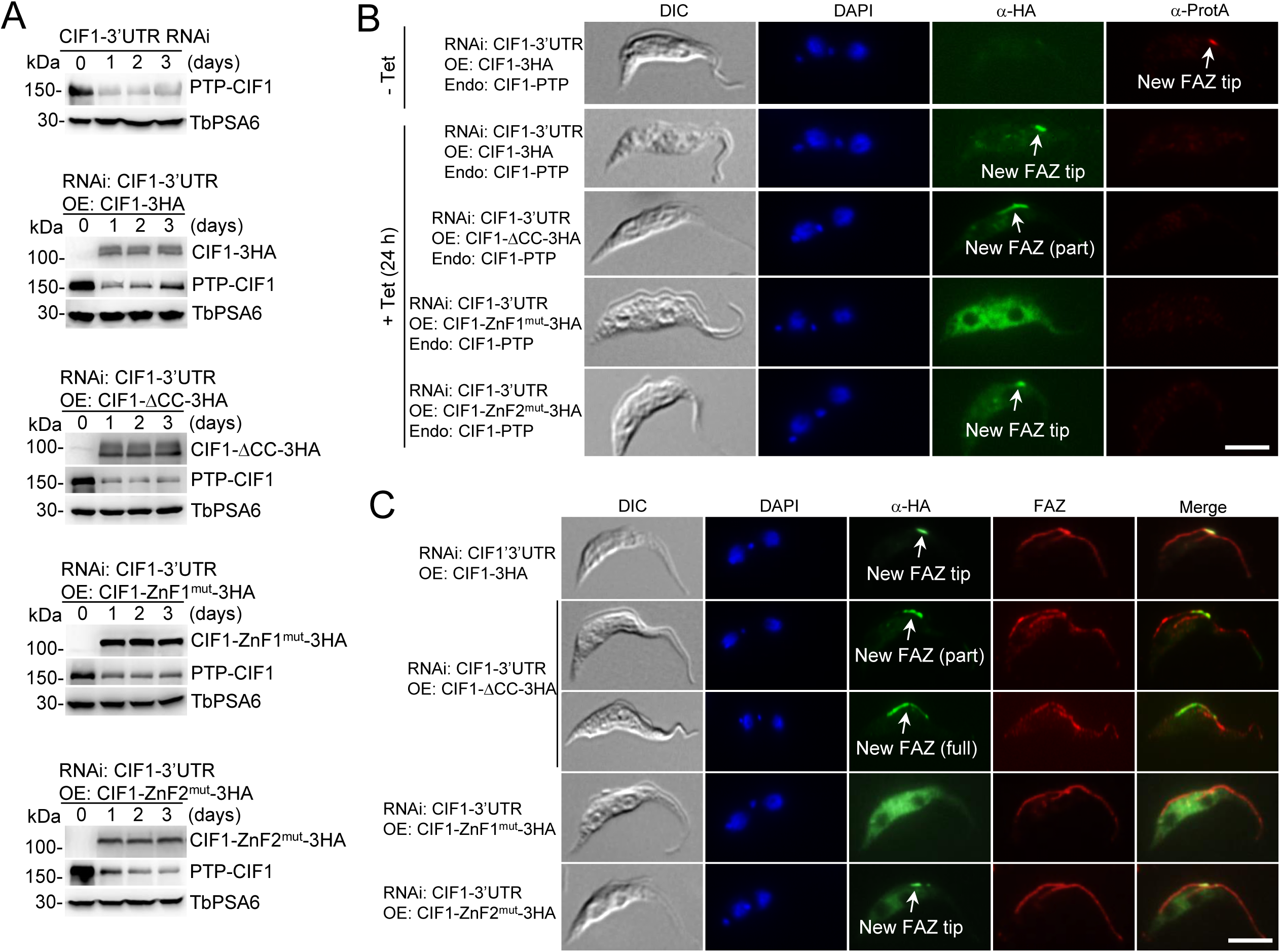
Complementation of CIF1-3’UTR RNAi by ectopically expressing CIF1 and its mutants. (**A**). Western blotting to detect the level of endogenous CIF1, which was tagged with a PTP epitope at the N-terminus, and the level of ectopically expressed CIF1 and its mutants, which were each tagged with a triple HA epitope at the C-terminus. Cells were incubated with 1.0 μg/ml tetracycline, and time-course samples were collected for western blotting with anti-Protein A pAb and anti-HA mAb. TbPSA6 level served as the loading control. (**B**). Immunofluorescence microscopy to detect the level of endogenous PTP-tagged CIF1 and ectopically expressed 3HA-tagged CIF1 and its mutants. Non-induced control and tetracycline-induced cells were co-immunostained with anti-Protein A pAb and FITC-conjugated anti-HA mAb, and counterstained with DAPI for nucleus and kinetoplast. Scale bar: 5 μm. (**C**). Immunofluorescence microscopy to monitor the localization of ectopically expressed CIF1 and its mutant in CIF1-3’UTR RNAi cells. Cells were co-immunostained with FITC-conjugated anti-HA mAb and anti-CC2D pAb, which labels the FAZ filament. Scale bar: 5 μm.

We next investigated whether ectopic CIF1 and its mutants were able to complement CIF1-3’UTR RNAi. Expression of wild-type CIF1 in the CIF1-3’UTR RNAi cell line restored cell growth (Fig. 6A), demonstrating the complementation of CIF1 deficiency by the ectopic CIF1 protein. However, expression of CIF1-ΔCC in CIF1-3’UTR RNAi cells failed to restore cell growth (Fig. 6A), indicating that the coiled-coil motif is important for CIF1 function. Given that the coiled-coil motif is required for restricting CIF1 to the new FAZ tip (Figs. 4B and 5B, C), these results suggest that localization of CIF1 to the anterior tip of the new FAZ is essential for CIF1 to execute its functions.

**Figure 6.**
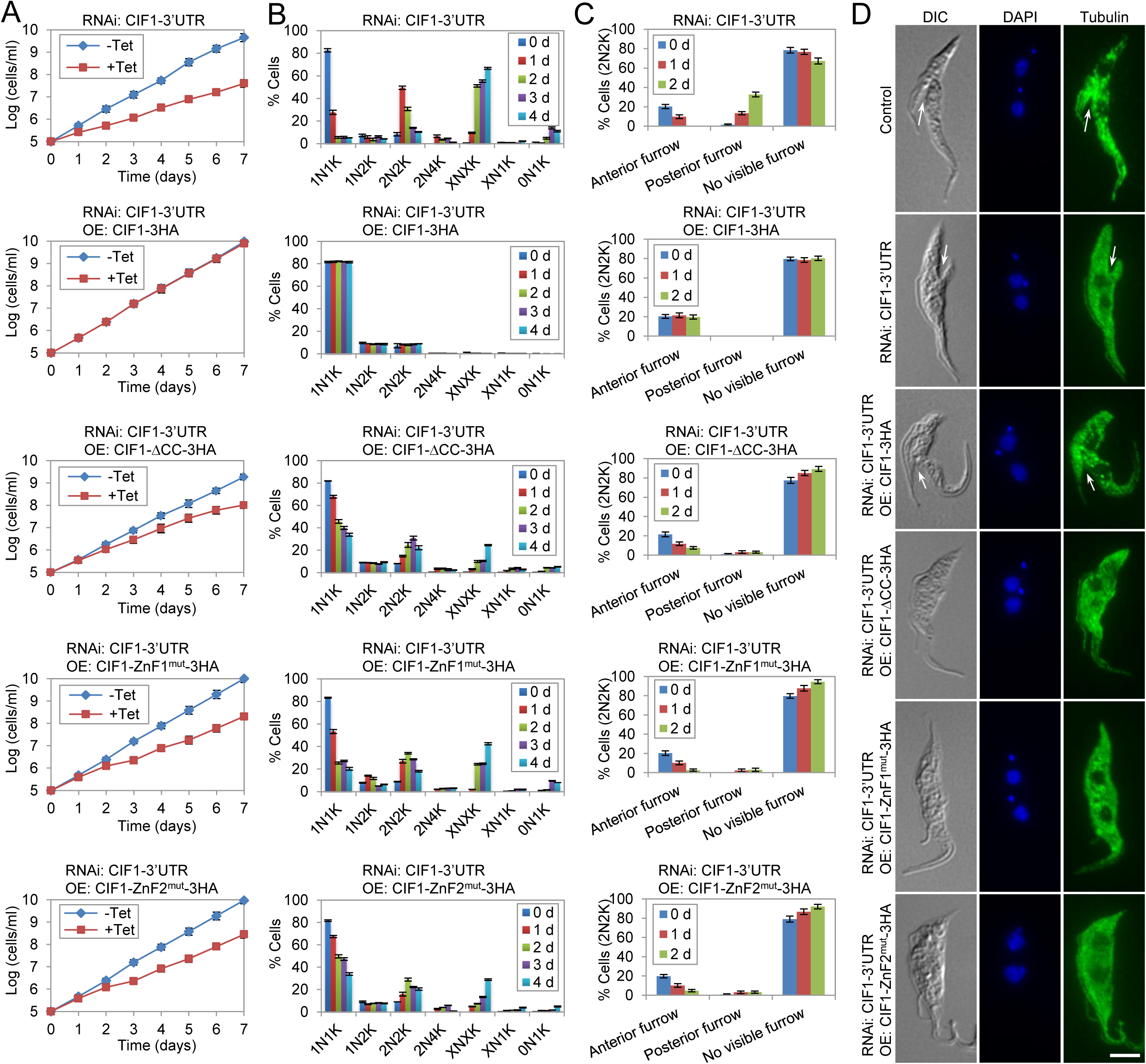
The structural motifs in CIF1 are required for CIF1 function in cytokinesis. (**A**). RNAi complementation experiments to investigate the requirement of the structural motifs in CIF1 for cell proliferation. Shown are the growth curves of CIF1-3’UTR RNAi cell line and CIF1-3’UTR RNAi cell lines expressing wild-type CIF1 or CIF1 mutants. Error bars indicate S.D. from three independent experiments. (**B**). Effect of mutation of the structural motifs in CIF1 on cell cycle progression. CIF1-3’UTR RNAi cell line and CIF1-3’UTR RNAi cell lines expressing wild-type CIF1 or CIF1 mutants were induced with tetracycline, and time-course samples were stained with DAPI for counting the cells with different numbers of nucleus and kinetoplast. A total of 300 cells were counted for each time point, and error bars indicate S.D. from three independent experiments. (**C**). Effect of mutation of the structural motifs in CIF1 on cleavage furrow ingression. CIF1-3’UTR RNAi cell line and CIF1-3’UTR RNAi cell lines expressing CIF1 or its mutants were induced with tetracycline for 2 days. Non-induced control and tetracycline-induced samples were fixed, stained with DAPI, and examined under a fluorescent microscope to examine the cleavage furrow in 2N2K cells. A total of 200 cells were counted for each time point, and error bars indicate S.D. from three independent experiments. (**D**). Visualization of the cleavage furrow in control, CIF1-3’UTR RNAi, and RNAi complementation cell lines. Cells were immunostained with anti-α-tubulin mAb and counterstained with DAPI for nuclear and kinetoplast DNA. The white arrow pointed to the anterior furrow in a control cell and a CIF1-complemented CIF1-3’UTR RNAi cell and the posterior furrow in a CIF1-3’UTR RNAi cell. Scale bar: 5 μm.

Due to the mis-localization of CIF1-ZnF1^mut^ to the cytosol (Fig. 5B, C), expression of CIF1-ZnF1^mut^ in CIF1-3’UTR RNAi cells failed to complement the loss of endogenous CIF1 protein, causing a growth defect (Fig. 6A), demonstrating that ZnF1 is essential for CIF1 function. Surprisingly, expression of CIF1-ZnF2^mut^ was unable to restore cell growth (Fig. 6A), indicating that CIF1-ZnF2^mut^ was incapable of rescuing CIF1-3’UTR RNAi, despite being able to localize to the new FAZ tip (Fig. 5B, C). These results suggest that ZnF2 is also required for CIF1 function.

### Effect of the mutation of CIF1 structural motifs on cytokinesis initiation

We next investigated the effect of the expression of CIF1 mutants in CIF1-3’UTR RNAi cells on cytokinesis initiation. As the control, we first examined the cytokinesis defects of CIF1-3’UTR RNAi cells. Quantification of cells with different numbers of nucleus and kinetoplast showed that depletion of CIF1 caused a significant increase of 2N2K cells from ∼10% to ∼50% of the total population after 2 days of RNAi induction and then an accumulation of XNXK (X>2) cells to ∼65% of the total population after RNAi induction for 4 days (Fig. 6B). Quantification of the number of 2N2K cells with a visible cleavage furrow showed that upon CIF1 RNAi the 2N2K cells with a visible anterior cleavage furrow decreased from ∼20% to less than 1%, whereas the 2N2K cells with a visible posterior cleavage furrow emerged to ∼30% of the total 2N2K population (Fig. 6C). These cytokinesis defects are similar to the CIF1 RNAi cell line through targeting the CIF1 coding sequence (5), further confirming that CIF1 is required for cytokinesis initiation from the anterior cell end and that depletion of CIF1 caused cleavage furrow ingression from the posterior cell end.

Consistent with the restoration of cell growth, ectopic expression of CIF1 in CIF1-3’UTR RNAi cells did not affect cell cycle progression (Fig. 6B) and cleavage furrow ingression (Fig. 6C, D). However, ectopic expression of CIF1-ΔCC in CIF1-3’UTR RNAi cells caused an increase of 2N2K cells from ∼10% to ∼25% of the total population after tetracycline induction for 3 days and an accumulation of XNXK (X>2) cells to ∼25% of the total population after induction for 4 days (Fig. 6B). Moreover, the 2N2K cells with a visible anterior cleavage furrow were reduced from ∼20% to ∼7% (Fig. 6C, D), indicating a defect in cytokinesis initiation from the anterior cell end. Intriguingly, the 2N2K cells with a posterior cleavage furrow were not significantly increased as in CIF1-3’UTR RNAi cells (Fig. 6C, D). Ectopic expression of CIF1-ZnF1^mut^ or CIF1-ZNF2^mut^ in CIF1-3’UTR RNAi cells both resulted in a significant increase of 2N2K cells from ∼10% to ∼34% or 29% after tetracycline induction for 2 days and the accumulation of XNXK cells to ∼42% or 29% of the total population after tetracycline induction for 4 days (Fig. 6B). Moreover, the 2N2K cells with a visible anterior cleavage furrow were significantly reduced from ∼20% to ∼2-5% after tetracycline induction for 2 days, and the 2N2K cells with a posterior cleavage furrow were not significantly increased in both cell lines (Fig. 6C, D). These results indicated that both zinc-finger motifs are required for CIF1 function in cytokinesis initiation. These results also suggest that the presence of non-functional CIF1 protein, such as CIF1-ΔCC, CIF1-ZnF1^mut^, or CIF1-ZnF2^mut^, likely suppressed cleavage furrow ingression from the posterior cell end in CIF1-3’UTR RNAi cells.

### Identification of CIF1 structural motifs required for CIF2 localization and stability

The identification of the structural motifs required for CIF1-CIF2 interaction promoted us to investigate whether interaction with CIF1 is required for CIF2 localization to the new FAZ tip. Immunofluorescence microscopy showed that in CIF1-3’UTR RNAi cells expressing wild-type CIF1, PTP-tagged CIF2 was localized to the new FAZ tip and co-localized with CIF1, which was tagged with a C-terminal triple HA epitope, in almost all the S-phase cells examined (Fig. 7A, B). In CIF1-3’UTR cells expressing CIF1-ΔCC, PTP-CIF2 was detected either at the anterior one-third length of the new FAZ or along the full length of the new FAZ, where it co-localized with CIF1-ΔCC, in almost all of the S-phase cells (1N1K cells with an elongated kinetoplast and a short new FAZ) examined (Fig. 7A, B). In CIF1-3’UTR RNAi cells expressing CIF1-ZnF1^mut^, however, PTP-tagged CIF2 was not detected at the new FAZ tip, but in the cytosol, in all the S-phase cells examined (Fig. 7A, B). In these cells, CIF1-ZnF1^mut^ was also detected in the cytosol (Fig. 7A). In CIF1-3’UTR RNAi cells expressing CIF1-ZnF2^mut^, PTP-CIF2 was detected in the cytosol, despite that CIF1-ZnF2^mut^ was still localized to the new FAZ tip in all S-phase cells examined (Fig. 7A, B). Given that CIF1-ZnF2^mut^ did not interact with CIF2 (Fig. 2F), these results suggest that CIF2 localization to the new FAZ tip depends on its interaction with CIF1.

**Figure 7.**
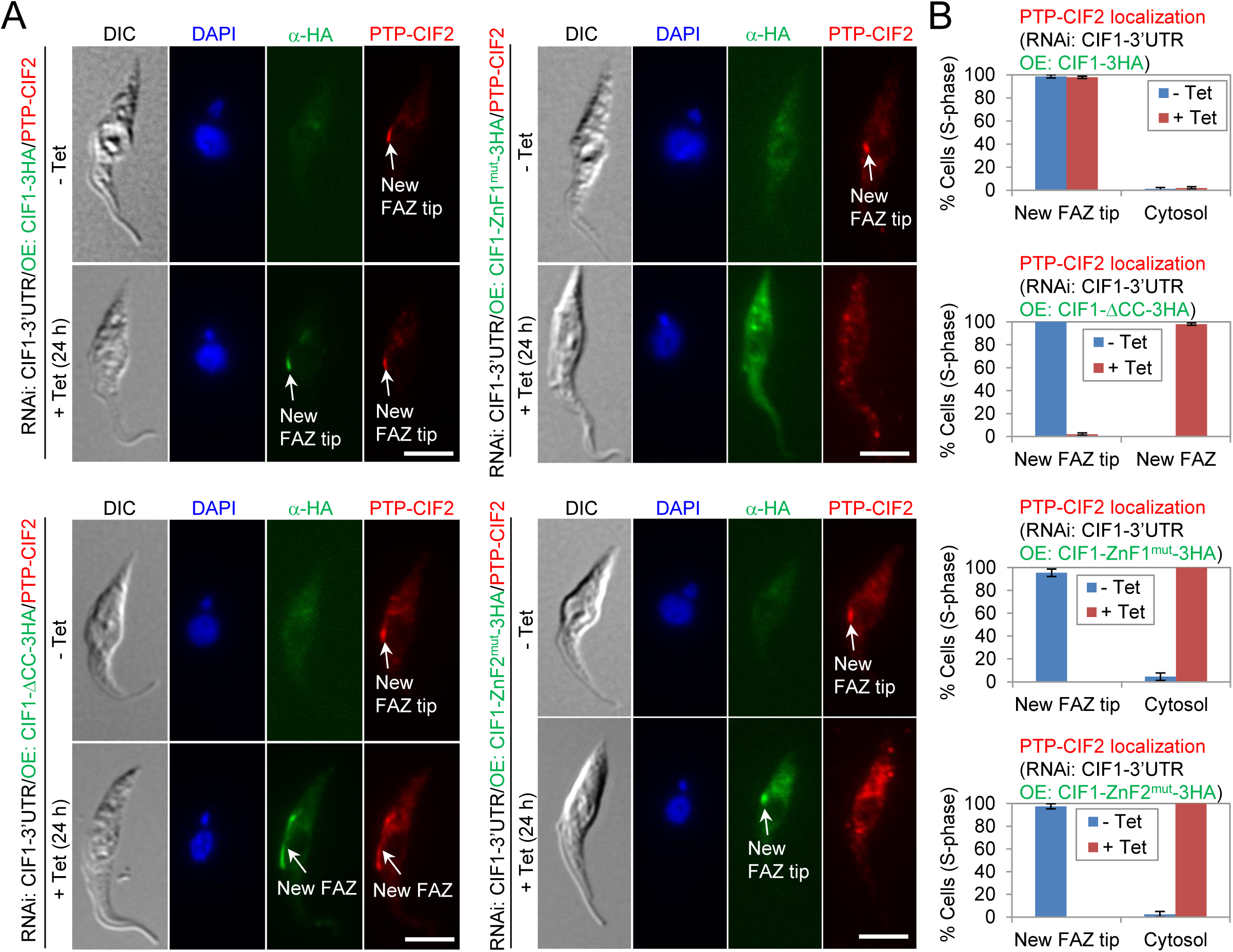
Requirement of the structural motifs in CIF1 for CIF2 localization. (**A**). Immunofluorescence microscopy to detect the subcellular localization of CIF2-PTP in CIF1-3’UTR RNAi cell lines expressing CIF1-3HA, CIF1-ΔCC-3HA, CIF1-ZnF1^mut^-3HA, or CIF1-ZnF2^mut^-3HA. Non-induced control and tetracycline-induced cells (24 h) were immunostained with anti-Protein A pAb to detect CIF2-PTP and with FITC-conjugated anti-HA mAb to detect 3HA-tagged wild-type and mutant CIF1 proteins. Scale bar: 5 μm. (**B**). Quantification of CIF2 localization in control and CIF1-3’UTR RNAi cell lines expressing wild-type and mutant CIF1 proteins. A total of 200 S-phase (1N1K cells with an elongated kinetoplast and a short new FAZ) cells were counted for each time point, and results were presented as mean percentage ± SD (*n*=3).

We next investigated the effect of CIF1 mutation on the stability of CIF2 protein. In CIF1-3’UTR RNAi cells, PTP-tagged CIF2 was gradually decreased upon RNAi induction (Fig. 8A, B). In CIF1-3’UTR RNAi cells expressing wild-type CIF1, CIF2 level was not affected (Fig. 8A, B), consistent with the rescue of CIF1-3’UTR RNAi phenotype by ectopic expression of CIF1. In contrast, in CIF1-3’UTR RNAi cells expressing CIF1-ΔCC, CIF2 level was instead increased ∼35% after tetracycline induction for 72 h (Fig. 8A, B). In CIF1-3’UTR RNAi cells expressing either CIF1-ZNF1^mut^ or CIF1-ZnF2^mut^, CIF2 protein level gradually decreased (Fig. 8A, B). Given that wild-type CIF1 and CIF1-ΔCC, but not CIF1-ZnF1^mut^ and CIF1-ZnF2^mut^ (Fig. 2F), interact with CIF2 *in vivo* in trypanosome, these results suggest that the interaction of CIF2 with CIF1 is required to maintain CIF2 protein stability.

**Figure 8.**
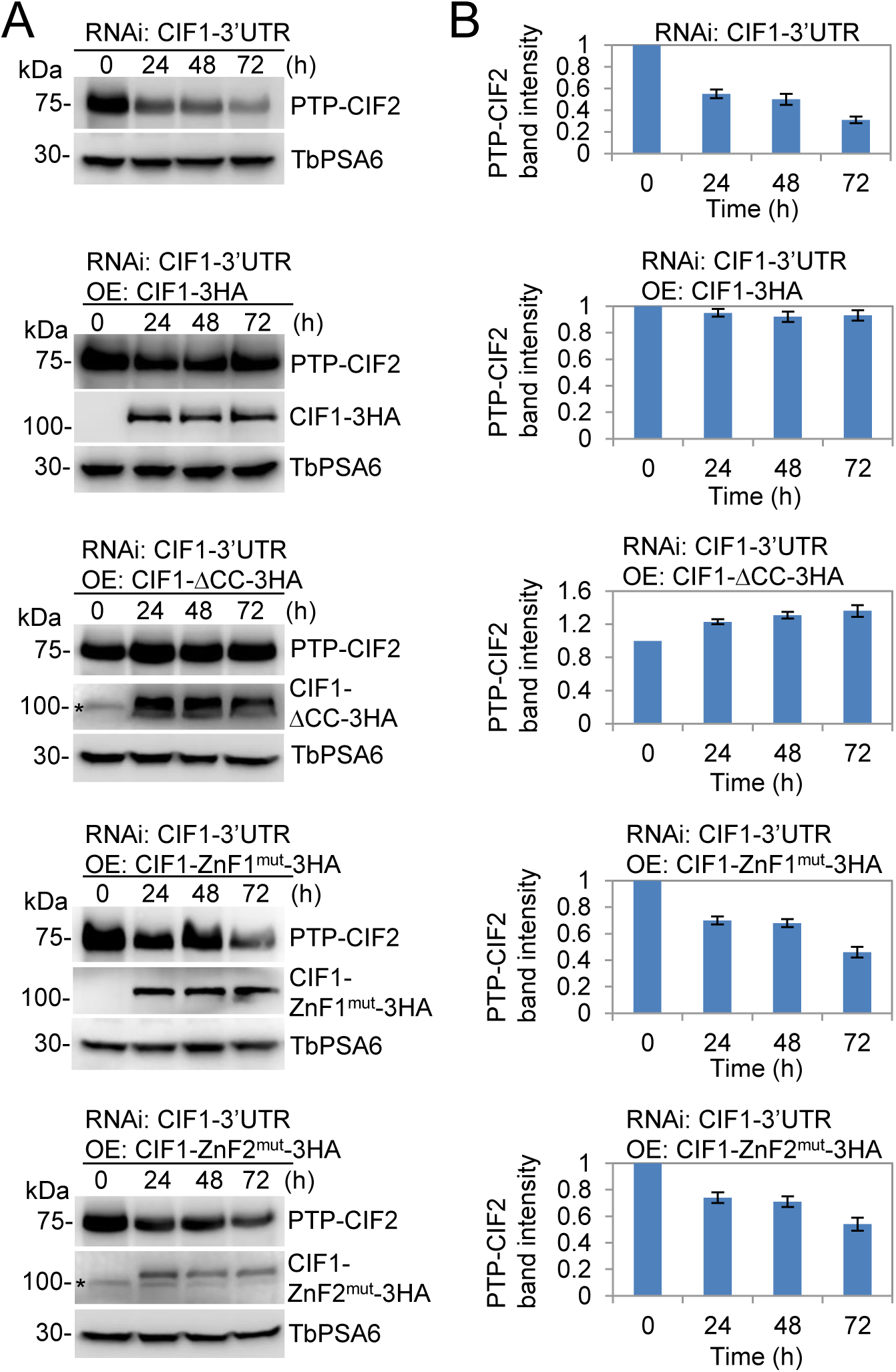
CIF1 structural motifs required for maintaining CIF2 protein stability. (**A**). Western blotting to examine the level of CIF2, which was tagged with a C-terminal PTP epitope from one of its endogenous loci, and the level of CIF1 and its mutants, which were tagged with a C-terminal triple HA epitope and ectopically overexpressed in cells containing the CIF1-3’UTR RNAi construct. Cells were induced with tetracycline for 72 h, and time-course samples were collected for western blotting with anti-Protein A antibody and anti-HA antibody. TbPSA6 served as the loading control. The asterisks indicate a non-specific band detected by anti-HA antibody. (**B**). Quantification of the level of CIF2 protein in CIF1-3’UTR RNAi cell line and CIF1-3’UTR RNAi cell lines expressing CIF1 and its mutants. CIF2 band intensity was determined by ImageJ, and normalized with the intensity of TbPSA6. Error bars indicate S.D. calculated from three independent experiments.

## Discussion

Cytokinesis initiation in trypanosomes requires the CIF1-CIF2 protein complex that acts at the anterior tip of the new FAZ during the S phase of the cell cycle (5,21). Through bioinformatics analyses and homology modeling, we identified a coiled-coil motif and two CCHC-type zinc-finger motifs in CIF1 and a calmodulin-like domain consisting of four EF-hand motifs in CIF2 (Fig. 1). All these motifs, which are present in many eukaryotic proteins, are known to be involved in protein-protein interactions (28-31), and interactions between calmodulin and its binding proteins can be either calcium-dependent or calcium-independent (32,33). Indeed, through yeast two-hybrid, size exclusion chromatography, and co-immunoprecipitation, we identified the two ZnF motifs in CIF1 and the calmodulin-like domain in CIF2 as the domains involved in CIF1-CIF2 interaction (Figs. 2 and 3). We also showed that interaction between CIF1 and CIF2 is not dependent on calcium (Fig. 2D). This finding and the observation that the calmodulin-like domain in CIF2 lacks the canonical Ca^2+^ binding sites (Fig. 1B) raise the question of whether CIF2 is capable of binding to calcium.

The RNAi complementation experiments through ectopically expressing various CIF1 mutants in CIF1-deficient cell lines allowed us to investigate the requirement of the structural motifs for the subcellular localization and the physiological functions of CIF1. The finding that the CIF1-ΔCC ectopic expression/CIF1-3’UTR RNAi cell line, the CIF1-ZnF1^mut^ ectopic expression/CIF1-3’UTR RNAi cell line, and the CIF1-ZnF2^mut^ ectopic expression/CIF1-3’UTR RNAi cell line all exhibited a growth defect (Fig. 6A) and a cytokinesis defect (Fig. 6B, C) demonstrated that the coiled-coil motif and the two zinc-finger motifs are all required for CIF1 function in cytokinesis. Notably, the three CIF1 mutants displayed distinct subcellular localizations (Fig. 5B, C), suggesting that the cytokinesis defects observed in the three cell lines might be due to different mechanisms. The fact that CIF1-ΔCC was spread over to the anterior one-third length or the full length of the new FAZ (Fig. 5B,C) and was not able to complement CIF1 deficiency (Fig. 6) indicated that the coiled-coil motif is required for restricting CIF1 to the new FAZ tip so that CIF1 is able to execute its function. The underlying mechanism for this function of the coiled-coil motif is not clear, but presumably it could be due to the disruption of CIF1 interaction with a yet unknown FAZ tip-localizing protein that confines CIF1 at the new FAZ tip. This observation is similar to the effect of TbSAS-4 RNAi on the localization of several TbSAS-4-associated proteins, which were spread over to the anterior one-third length of the new FAZ in TbSAS-4 RNAi cell lines (34). Another possible mechanism underlying the cytokinesis defect caused by CIF1-ΔCC could be the increase in CIF2 protein level (Fig. 8), as overexpression of CIF2 was previously shown to disrupt cytokinesis (21). While the effect of ZnF1 mutation on CIF1 function (Fig. 6) is consistent with its mis-localization to the cytosol (Fig. 5B, C), mutation of ZnF2 disrupted CIF1 function but not localization (Figs. 5 and 6). This interesting result indicated that despite being able to localize to the new FAZ tip, CIF1-ZnF2^mut^ lost its function in cytokinesis, which in part could be attributed to the failure of targeting CIF2 to the new FAZ tip (Fig. 7) and the degradation of CIF2 (Fig. 8).

Our results demonstrated that CIF2 localization to the new FAZ tip depends on its interaction with CIF1. First, CIF2 mutants lacking the CIF1-interacting EF-hand motifs failed to localize to the new FAZ tip (Fig. 4D). Secondly, CIF2 in cells expressing CIF1 zinc-finger mutants that do not bind to CIF2 failed to localize to the new FAZ tip (Fig. 7). Finally, CIF2 in cells expressing CIF1-ΔCC mutant that still binds to CIF2 was spread over, together with CIF1-ΔCC, to the anterior one-third length or the full-length of the new FAZ (Fig. 7). Moreover, our results also demonstrated that CIF2 protein stability depends on its interaction with CIF1. CIF2 in CIF1-3’UTR RNAi cells expressing CIF1 zinc-finger mutants were gradually destabilized, whereas CIF2 in CIF1-3’UTR RNAi cells expressing CIF1-ΔCC was instead stabilized (Fig. 8). Together, these results suggest that CIF1 acts upstream of CIF2 by targeting CIF2 to the new FAZ tip during S phase.

In summary, through yeast two-hybrid and co-immunoprecipitation assays, we have determined the structural motifs required for CIF1-CIF2 complex formation, and through RNAi complementation experiments, we have determined the structural motifs required for CIF1 localization and function and demonstrated that CIF1 targets CIF2 to the new FAZ tip and maintains CIF2 stability. These results provided structural insights into the functional interplay between CIF1 and CIF2 and determined the order of action between the two proteins in the cytokinesis regulatory pathway.

## Experimental Procedures

### Generation of CIF1 and CIF2 mutants and yeast two-hybrid assays

To test the interaction between CIF1 and CIF2 by directional yeast two-hybrid assay, full-length coding sequence of CIF1 and CIF2 was cloned between the *EcoR*I and *BamH*I sites in pGAD and pGBK vectors (Takara Bio. USA). To generate the coiled-coil motif deletion mutant of CIF1 (CIF1-ΔCC), the coding sequences flanking the DNA sequence (nucleotides 363-813) that encodes the coiled-coil motif were fused and cloned into the pGBK vector. To generate the zinc-finger motif #1 mutant (CIF1-ZnF1^mut^), the putative zinc ion-coordinating residues Cys693, Cys696, His708, and Cys712 in ZnF1 were all mutated to alanine. Similarly, the four putative zinc ion-coordinating residues Cys769, Cys772, His784, and Cys788 in ZnF2 were mutated to alanine to generate the zinc-finger motif #2 mutant (CIF1-ZnF2^mut^). CIF1-ZnF1&2^mut^ was generated by mutating the putative zinc ion-coordinating residues in both zinc-finger motifs to alanine. These mutants were all cloned into the pGBK vector. To generate CIF2 EF-hand deletion mutants, CIF2 coding sequence without the sequence encoding the first EF-hand motif (EF1,a.a 1-46), the first two EF-hand motifs (EF1-2, a.a. 1-82), the first three EF-hand motifs (EF1-3, a.a. 1-119), or all four EF-hand motifs (EF1-4, a.a. 1-160) was cloned into the pGAD vector (see Table S1 for the sequences of all primers used in this study)

The pGBK plasmids were transformed into yeast stain Y187 (mating type α), and the pGAD plasmids were transformed into yeast strain AH109 (mating type a). Yeast mating was carried out by mixing the Y187 and AH109 strains and incubating the mixture at 30 ^o^C according to our published procedures (35,36). The diploid yeast strains were spotted onto SD plates lacking Leu and Trp (2DO, double drop-out, control) to select the presence of both the pGBKT7 and the pGADT7 plasmids. The diploid yeast strains were also spotted onto SD plates lacking Leu, Trp, and His (3DO, triple drop-out, medium stringency), or lacking Leu, Trp, His, and Ade (4DO, quadruple drop-out, high stringency) for detection of the interaction between the pray and bait proteins under medium stringency and high stringency conditions, respectively (for details, refer to the Matchmaker® yeast two-hybrid system manual from Takara Bio. USA).

### Purification of recombinant CIF1-CTD and CIF2-NTD and size exclusion chromatography

To generate constructs of CIF1-CTD and CIF2-NTD for co-expression in *E. coli*, we cloned the DNA sequence encoding CIF1-CTD (a.a. 667-804) and CIF2-NTD (a.a. 1-160) into the *Nde*I/*BamH*I sites of pET29a and the *Nde*I/*Bgl*II sites of pET15b (Novagen), respectively. The pET15b vector provides a thrombin-cleavable N-terminal 6×His tag, and is compatible with pET29a for protein co-expression in *E. coli*. Additionally, CIF1-CTD and CIF2-NTD were respectively cloned into the modified SUMO-pET15b and SUMO-pET28a vectors, which both provide a similar 6×His-SUMO tag at the N-terminus of the target proteins.

For co-expression, different combinations of the above constructs were used to co-transform competent *E. coli* BL21(DE3) cells. Bacterial cells were grown in the LB medium containing both 50 μg/ml ampicillin and 35 μg/ml kanamycin at 37 °C until their OD_600_ reached 0.6-1.0 (approximately 2-3 h). Protein expression was induced with 250 μM of isopropyl-beta-D-thiogalactopyranoside (IPTG) at 18°C overnight. Cells were harvested by centrifugation (6,000×*g*, 15 min, 4°C). Cell pellets were resuspended in pre-chilled lysis buffer containing 20 mM Tris-HCl (pH 8.0), 100 mM NaCl, 20 mM imidazole, 10 mM beta-mercaptoethanol, and 5% (v/v) glycerol. Cells were lysed using the EmulsiFlex-C3 homogenizer (Avestin). Cell debris was removed by centrifugation (25,000×g, 40 min, 4°C), and the supernatant was filtered through a 0.45 μm pore size filter and loaded onto a 5-ml Ni-HiTrap column (GE Healthcare) pre-equilibrated in the same lysis buffer. After washing with 5 column volumes (cv) of lysis buffer, bound proteins were eluted by a linear gradient concentration of imidazole (20 to 600 mM) in the lysis buffer over 10 × cv. In the case of CIF2-NTD in pET15b, the N-terminal 6×His tag was removed by incubation with ∼5% (w/w) of thrombin (4°C, overnight). Co-purified proteins were subjected to size exclusion chromatographic analyses using the Superdex S-200 16/60 column (GE Healthcare) pre-equilibrated with the running buffer containing 20 mM Tris-HCl (pH 8.0), 100 mM NaCl, and 5% (v/v) glycerol. Peak fractions from each elution were finally examined on a 15% (w/v) SDS-PAGE gel and stained with Coomassie blue.

### Trypanosome cell culture and RNAi

The procyclic *T. brucei* 29-13 strain (37) was cultured in SDM-79 medium supplemented with 10% heat-inactivated fetal bovine serum (Sigma-Aldrich), 15μg/ml G418, and 50 μg/ml hygromycin at 27^o^C.

To generate CIF1-3’UTR RNAi cell line, a 500-bp DNA fragment from the 3’UTR of CIF1, which does not overlap with the downstream gene, was cloned into the *Hind*III/*Xho*I sites of the pZJM-PAC vector, which was modified from the original pZJM vector (38) by replacing the phleomycin resistance gene with puromycin resistance gene. To generate CIF2-3’UTR RNAi cell line, a 500-bp DNA fragment from the 3’UTR of CIF2 was similarly cloned into the pZJM-PAC vector. The pZJM-CIF1-3’UTR-PAC and pZJM-CIF2-3’UTR-PAC plasmids were each linearized with NotI and transfected into the 29-13 strain by electroporation. Transfectants were selected with 1.0 μg/ml puromycin and cloned by limiting dilution in a 96-well plate. RNAi was induced with 1.0 μg/ml tetracycline, and cell growth was monitored daily by counting the cells with a hemacytometer. Three clonal cell lines for each RNAi strain were analyzed, which showed almost identical phenotypes, and results from one clonal cell lines were presented.

### RNAi complementation

Full-length CIF1, CIF1-ΔCC, CIF1-ZnF1^mut^, and CIF1-ZnF2^mut^ were each cloned into the *Xho*I/*Afl*II sites of the pLew100-3HA-BLE vector. The resulting plasmids were linearized with *Not*I and transfected into CIF1-3’UTR RNAi cell line. Similarly, full-length CIF2 was also cloned into the pLew100-3HA-BLE vector, and the resulting plasmid was linearized with *Not*I and transfected into CIF2-3’UTR RNAi cell line. Transfectants were selected under 2.5μg/ml phleomycin in addition to 1.0 μg/ml puromycin, 15 μg/ml G418 and 50 μg/ml hygromycin B, and then cloned by limiting dilution in a 96-well plate. Cells were induced with 1.0 μg/ml tetracycline, and cell growth was monitored daily by counting the cells under a light microscope with a hemacytometer. Three clonal cell lines for each of the RNAi complementation strains were analyzed.

In situ epitope tagging of CIF1 and CIF2*-*For tagging of CIF1 and CIF2 from one of their endogenous loci, the PCR-based tagging method (39) was carried out. CIF1 was tagged with a PTP epitope at the N-terminus in CIF1-3’UTR RNAi cell line and CIF1-3’UTR RNAi cell lines expressing 3HA-tagged CIF1 or CIF1 mutants. Similarly, CIF2 was tagged with an N-terminal PTP epitope in CIF1-3’UTR RNAi cell line and CIF1-3’UTR RNAi cell lines expressing 3HA-tagged CIF1 or CIF1 mutants. Transfectants were selected with 10 μg/ml blasticidin and cloned by limiting dilution in a 96-well plate.

### Ectopic expression of wild-type and mutant CIF1 and CIF2

To ectopically express CIF1, CIF2, and their respective mutants, the pLew100-3HA-BLE plasmids containing wild-type and mutant CIF1 and CIF2 that were constructed above for RNAi complementation were linearized with *Not*I and then transfected into the 29-13 strain. Transfectants were selected with 2.5 μg/ml phleomycin, and cloned by limiting dilution. Cells were incubated with 0.1 μg/ml tetracycline to induce the expression of 3HA-tagged CIF1, CIF2, and their respective mutants. Three clonal cell lines for each of the overexpression strains were analyzed.

### Immunoprecipitation and western blotting

Co-Immunoprecipitation was carried out essentially as described previously (21,36). Briefly, trypanosome cells (10^7^) were lysed in 1 ml lysis buffer (25 mM Tris-HCl, pH7.6, 150 mM NaCl, 1 mM DTT, 10% Nonidet P-40, and protease inhibitor cocktail) for 30 minutes on ice. Cleared cell lysate was incubated with 20 μl settled IgG beads (GE HealthCare) for 1 h at 4^o^C. The beads were then washed 6 times with the lysis buffer, and bound proteins were eluted with 10% SDS. For anti-HA immunoprecipitation, cells were similarly lysed and cleared by centrifugation. Cleared lysate was incubated with 20 μl settled EZview^TM^ Red anti-HA affinity gel (Sigma-Aldrich) for 1 h at 4^o^C. The beads were washed 6 times with the lysis buffer, and bound proteins were eluted with 10% SDS. Immunoprecipitated proteins were separated by SDS-PAGE, transferred onto a PVDF membrane, and immunoblotted with anti-HA mAb (clone HA-7, catalog #: H9658, Sigma-Aldrich, 1:2,000 dilution) to detect 3HA-tagged CIF1 and its various mutants and with anti-Protein A pAb (anti-ProtA) (catalog #: P3775, Sigma-Aldrich, 1:1,000 dilution) to detect PTP-tagged CIF2. As the controls, cells expressing CIF1-3HA only or PTP-CIF2 only were used for immunoprecipitation. Western blot images were acquired using the FluorChem® Q imaging system (ProteinSimple, San Jose, CA), which allows quantitative imaging without oversaturating in a single exposure.

To test whether CIF1-CIF2 interaction is dependent on calcium, cells expressing CIF1-3HA and PTP-CIF2 were incubated for 24 h with 10μM BAPTA-AM (Sigma-Aldrich), which is a membrane permeable calcium chelator that has been used in *T. brucei* recently (27), and then lysed for co-immunoprecipitation.

### Immunofluorescence microscopy

Cells were attached to the coverslips for 30 min, fixed with cold methanol (-20^o^C) for 30 min, and rehydrated with PBS for 10 min. Cells were then blocked with 3% BSA in PBS for 1 h at room temperature, and incubated with FITC-conjugated anti-HA monoclonal antibody (clone HA-7, catalog # H7411, Sigma-Aldrich, 1:400 dilution,) for 1 h at room temperature. Cells on the coverslips were washed three times with the washing buffer (0.1% Triton X-100 in PBS) and then incubated with anti-Protein A polyclonal antibody (catalog # P3775, Sigma-Aldrich, 1:400 dilution) or anti-CC2D polyclonal antibody (1:1,000 dilution) (3) for 1 h at room temperature. After three washes with the washing buffer, cells were incubated with Cy3-conjugated anti-rabbit IgG (catalog # C2306, Sigma-Aldrich, 1:400 dilution) for 1 h at room temperature. For anti-tubulin immunostaining, cells were similarly treated on the coverslips and then incubated with anti-α-tubulin monoclonal antibody (1:2,000 dilution; Sigma-Aldrich for 1 h at room temperature. After three washes with the washing buffer, cells were then incubated with FITC-conjugated anti-mouse IgG (catalog # F0257, Sigma-Aldrich, 1:400 dilution) for 1 h at room temperature. After three washes with the washing buffer, slides were mounted with DAPI-containing VectaShield mounting medium (Vector Labs). Cells were imaged with an inverted fluorescence microscope (Olympus IX71) equipped with a cooled CCD camera (model Orca-ER, Hamamatsu) and a PlanApo N 60x1.42-NA lens, and acquired using the Slidebook 5 software.

## Acknowledgements

We are grateful to Dr. Cynthia Y. He of the National University of Singapore for providing the anti-CC2D antibody. This work was supported by the NIH R01 grant AI101437 to Z. L. and by the Austrian Science Fund (FWF) grant P28231-B28 to G. D.

